# Monitoring the bioenergetic state and cell lysis of *Bacillus subtilis* throughout the growth cycle

**DOI:** 10.1101/2025.08.21.671461

**Authors:** Maria Dakes Stavrakakis, Madeleine Humphrey, Tjeerd van Rij, Colin R. Harwood, Henrik Strahl

## Abstract

**Background:** The Gram-positive bacterium *Bacillus subtilis* is a rapidly growing and easily manipulated microbe with a long history of exploitation for the commercial production of industrial enzymes, high-value biochemicals, antibiotics and other secondary metabolites. More recently, it has been developed as a food additive and plant probiotic . It grows on cheap substrates and remains productive during extended batch-fed growth conditions. Extensive knowledge of its genetics, biochemistry and gene regulation has facilitated the use of metabolic engineering strategies to optimise substrate utilisation and product yield. As a free-living environmental organism, *B. subtilis* differentiates into subpopulations with distinct biological functions (e.g. sporulation, biofilm formation, antimicrobials production, etc). However, during fermentation, cellular differentiation processes can pose a challenge for its optimal biotechnological utilization, particularly when emerging subpopulations do not contribute to product biosynthesis. Here, we present robust assays that facilitate the analysis of two previously difficult-to-study population properties of *B. subtilis*: (i) the energization levels of individual cells within post-exponential but actively growing cultures and (ii) the extent of cell lysis that can occur under such conditions.

**Results:** Our findings reveal an unappreciated level of heterogeneity in cell energization within post-exponential cultures, and a surprisingly high degree of cell lysis in seemingly healthy, actively growing populations. These data provide insights and add to our understanding of the biological complexities and single-cell heterogeneities present in superficially simple bacterial clonal cultures. They establish robust and well-validated analytical tools with which to study the associated processes and provide a foundation for further optimizing *B. subtilis* as an industrial production host.

**Conclusions:** Considerable research efforts have been aimed at increasing the productivity of *B. subtilis* for industrial, medical and agricultural products. However, its ability to undergo physiological and morphological differentiation processes at high cell densities ultimately limits its productivity. Our research reveals how the resulting heterogeneity impacts the population-level energy status of individual cells in the culture and the surprisingly high extent of population-level cell lysis. The ability to monitor these processes provides tools for evaluating the impact of genetic and metabolic engineering strategies to improve productivity.

## Introduction

The Gram-positive endospore-forming bacterium *Bacillus subtilis* and its close relatives are important industrial organisms, responsible for the commercial production of a large variety of industrial enzymes (e.g. amylases, pectinases, proteases, cellulases), high-value biochemicals (e.g. vitamins, inositol, hyaluronic acid, N-acetylglucosamine, isobutanol), antibiotics and secondary metabolites (1) . In its natural environment, this so-called free-living single-celled organism nevertheless engages in a range of social behaviours that positively impacts on its long-term survival . In many cases, this involves the formation of complex communities, with both kith and kin, that can either be cooperative (beneficial) or antagonistic (competitive) (2). *B. subtilis* forms complex communities in which bet-hedging and social differentiation are important survival strategies (3, 4). Ultimately, this results in a minority of the population undergoing a morphological differentiation process to form dormant and highly temperature, desiccation, and chemically resistant endospores (5). Sporulation is a complex and energy-demanding process that requires significant remodelling of metabolic activities and cellular structures and, as a result, *B. subtilis* communities strive to delay the commitment to this differentiation pathway by seeking and utilising alternative nutrient sources (6). This is achieved by developing clonal, yet physiologically distinct cell types specialised in foraging (motility, swarming and chemotaxis), horizontal gene transfer (competence and transformation), utilisation of complex substrates (secretion of hydrolases), and production of antagonists to combat competitors (secondary metabolites and antimicrobial peptides) (6–9). This ability to differentiate into subpopulations with distinct phenotypes provides *B. subtilis* with a selective advantage in its native environments, the soil, rhizosphere and phylloplane. In industrial bioreactors, sporulation mutants prevent spore formation and contamination of the next fermentation in multipurpose plants, ensuring there are no viable cells present in the final product. However, such mutants do not eliminate processes that have evolved to delay sporulation.

In addition to the above-mentioned, well-defined differentiation pathways, *B. subtilis* exhibits autolytic behaviour upon de-energisation triggered by extreme nutrient-limiting conditions (10, 11). As a result, a proportion of the cell population becomes de-energised and undergoes cell lysis, releasing cell contents and nutrients that allow the remaining cell population to sustain metabolic activity and energisation (12) . In addition to relying on stochastic processes leading to de-energisation and the associated autolysis, cells in the early but still reversible stages of sporulation synthesise and secrete two proteins, the sporulation delaying protein (SdpC) and the sporulation killing factor (SkfA) that trigger lysis of other members of the community (13). This represents a form of active cannibalism that affects a proportion of non-sporulating cells, ultimately providing an additional nutrient source for the remaining cell population (14). This programmed cell lysis allows the population to delay or even avoid the induction of the energy-intensive differentiation pathway leading ultimately to sporulation.

While these complex differentiation processes and the associated population heterogeneities provide an advantage in its natural environments, they can represent a distinct disadvantage in industrial fed-batch bioreactor cultivation conditions, where optical densities of over 100 are commonly required (15). Both programmed and energy depletion-linked autolysis can reduce productivity and release intracellular products (e.g. nucleic acids, endotoxins, proteases), that are detrimental to the final product (16). This means that genetic and metabolic engineering strategies aimed at reducing cell lysis and optimising fermentation yield require effective assays to monitor variations in both the energetic status of individual cells in the population, and the extent of cell lysis, at different stages of growth.

At the single-cell level, bacterial viability and lysis are commonly monitored by so-called viability or Live/Dead assays (17), which find their use in a wide range of applications including microbiological quality control assessments of environmental or industrial samples and studies on the effectiveness and mode of action of antibiotics (18). Despite the arguably misleading commercial names, these types of assays do not report viability *per se* but utilise membrane-impermeable DNA-intercalating fluorescent dyes such as SYTOX Green and propidium iodide, which are excluded from bacterial cells with intact cell membranes (19, 20). When the membrane permeability barrier is compromised through membrane-active compounds or when cells undergo lysis, these dyes penetrate the cell and stain cellular DNA (21, 22). However, these assays cannot detect more subtle disturbances of the membrane permeability barrier function such as smaller-sized pores or increased ion conductivity and, indeed, are blind to metabolic inactivity unless associated with a severe membrane disruption or cell lysis . In contrast, voltage-sensitive fluorescent probes such as the carbocyanine dye, 3,3’-dipropylthiadicarbocyanine iodide (DiSC_3_(5)) can be used to assay cell membrane potential, which provides a more direct readout for the cell’s bioenergetic status and a more sensitive proxy for the state of the membrane permeability barrier function. Because of its cationic and hydrophobic characteristics, DiSC_3_(5) accumulates in polarised cells in a voltage-dependent manner. Upon membrane depolarisation, the dye is released from the cell, resulting in a reduced fluorescence signal. This voltage-dependent staining behaviour allows the membrane potential levels to be monitored at the single-cell level using flow cytometry or fluorescence microscopy (23, 24). The use of DiSC_3_(5) as a single-cell reporter in *B. subtilis* is well established (23, 25). However, due to cell density-related complexities in staining, and the maintenance of native membrane energisation during sample processing, the use of DiSC_3_(5) has been limited to early exponential growth phase or low cell density cultures, restricting comparisons only to cultures with similar cell densities. Consequently, there is a lack of reliable protocols for monitoring bacterial bioenergetic status using voltage-sensitive dyes at the single-cell level from more varied culture conditions.

To facilitate cell energetic status measurements during fermentation, we provide a systematic analysis of factors relevant to measuring *B. subtilis* membrane potential across a range of cell densities, thereby significantly expanding the applicability of DiSC_3_(5) as a sensitive proxy for cellular bioenergetic status. Furthermore, we have established how a combination of DiSC_3_(5) and SYTOX Green can be used to monitor cell energisation levels and cell lysis across the different growth phases of *B. subtilis*. This combination of methods provides insights into an unexpectedly high level of population heterogeneity in cell energisation and lysis that can occur despite seemingly healthy growth of the overall culture. While our study is limited to *B. subtilis*, we are confident that the established methods provide a solid foundation for developing similar assays in other microorganisms.

## Materials and methods

### Strains, media and growth conditions

The bacterial strains used in this study and their associated genotypes are listed in Table 1. Strain 168 (*trpC2*) is a tryptophan auxotroph due to a three-base pair deletion in *trpC*, encoding indol-3-glycerol-phosphate synthetase (Fig. S1). We generated a prototrophic variant of 168 to avoid the need to add tryptophan to the minimal salts culture medium. Previous attempts to generate such prototrophic strains of 168 have often used DNA from strain W23, which is only 94.6% identical to 168 and can introduce off-target nucleotide variations (26,27). To ensure the restoration of an authentic prototrophic version of *trpC*, we transformed strain 168 with a PCR-generated product of the *trpC* gene from strain NCiB3610^T^, amplified using oligos 5’-CTTCACAACCTCACTCCCTAAACAAAGC and 5’-GGAAGAGATCTATGCTTGAAAAAATCATCAAAC. The *trpC* gene of NCiB3610^T^ is identical to that of 168 except for the absence of a 3-bp deletion at nucleotides 423-5 (26). Following selection on minimal medium plates, transformant MDS23 was purified and its genome sequenced to confirm this strain carries the prototrophic *trpC* gene, while being otherwise genetically identical to the reference strain 168 (28) (Supplementary Figure S1).

**Table 1:** The list of bacterial strains used in this study. The list describes the name, genotype and origin of the strains. Antibiotic resistance markers: tetracycline (*tet*), spectinomycin (*spec*), neomycin (*neo*), kanamycin (*kan*), chloramphenicol (*cat*).

| Strain | Genotype | Construction or Reference |
| --- | --- | --- |
| *B. subtilis* 168 | *trpC2* | (28, 48, 49) |
| *B. subtilis* NCIB3610 | *trp*<sup>+</sup> | (50) |
| *B. subtilis* BKK03430 | *trpC2 nucA::kan* | (51) |
| *B. subtilis* BKK25750 | *trpC2 nucB::kan* | (51) |
| *B. subtilis* KS19 | *trpC2 lytABC::neo, lytD::tet, lytF::spec, lytE::cat* | (52) |
| *B. subtilis* MDS23 | *trp*<sup>+</sup> wild type (prototrophic derivative of 168) | this work |
| *B. subtilis* MDS29 | *trp*<sup>+</sup> *nucA::kan* | this work |
| *B. subtilis* MDS30 | *trp*<sup>+</sup> *nucB::kan* | this work |

Overnight cultures of *B. subtilis* were routinely grown in Lysogeny broth medium (LB; 0.5% (w/v) yeast extract, 1% (w/v) tryptone, 1% (w/v) NaCl), at 30°C and supplemented with 0.2% glucose to suppress sporulation. For the main experiments, strains were diluted 1:100 and grown at 37°C in modified Spizizen-minimal medium (mSMM: 0.2% (w/v) (NH_4_)_2_SO_4_, 0.6% (w/v) KH_2_PO_4_, 1.4% (w/v) K_2_HPO_4_, 0.12% (w/v) trisodium citrate (Na_3_C_6_H_5_O_7_.2H_2_O), 0.02% (w/v) MgSO_4._7H_2_O, 0.0011% (w/v) ammonium ferric citrate ((NH_4_)_5_Fe(C_6_H_4_O_7_)_2_), 0.5% (w/v) glucose, 0.02% (w/v) casamino acids). When growing the tryptophan auxotrophic *B. subtilis* strain 168, 0.002% (w/v) L-tryptophan was also added to the minimal medium. The autolysis experiments shown in Figure 1 and Supplementary Movie 1 were carried out in LB medium . In this article we use the widely avoid the misleading term ‘stationary growth phase’, replacing it with ‘post-exponential growth phase’ to indicate cultures that are growing, but no longer in the exponential growth phase.

**Figure 1:**
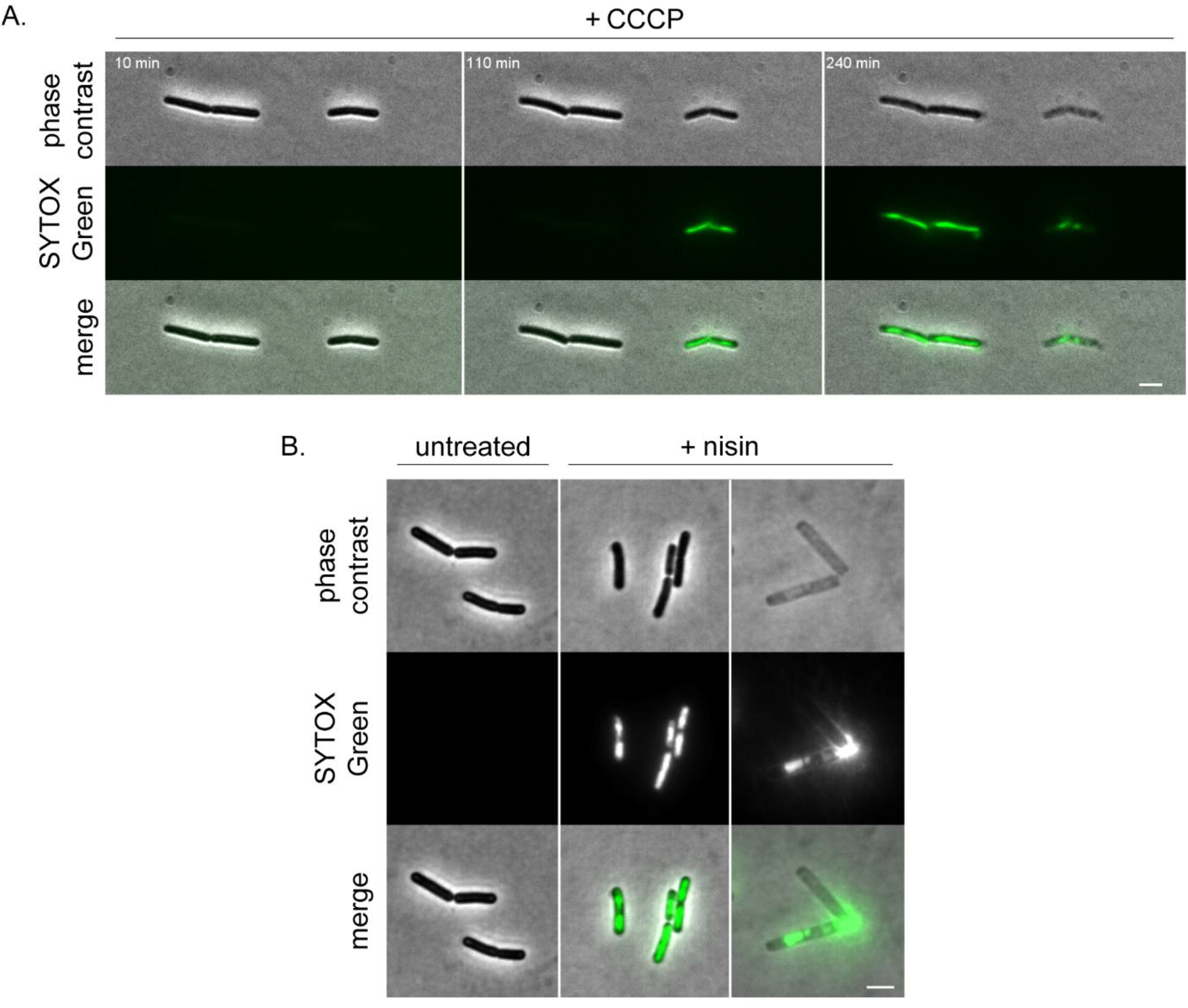
Autolysis of *B. subtilis* can be monitored with the membrane permeability-indicator SYTOX Green. **(A)** *B. subtilis* phase contrast and fluorescence time-lapse microscopy images of cells stained with 200 nM SYTOX Green in the presence of 100 μM of the proton uncoupler CCCP. For the full time-lapse experiment, see Movie S1. **(B)** *B. subtilis* phase contrast and fluorescence microscopy images of cells stained with 200 nM SYTOX Green in the absence (left panels) and presence (right panels) of the membrane-targeting, pore-forming lantibiotic nisin (10 μM for 5 min). The images are presented using identical contrast settings allowing a direct comparison between phase dark/SYTOX negative cells with intact membranes (left panels), phase dark/SYTOX Green positive cells with permeabilised membranes (middle panels) and extensively lysed phase light/ SYTOX Green positive cells (right panels). Scale bar: 3 μm. Strain used: (a) *B. subtilis* wild type (168) and (b) *B. subtilis* prototroph (MDS23).

### Fluorescence microscopy

For fluorescence microscopy, the cells were incubated with either 1μM of the voltage-sensitive dye DiSC_3_(5) dissolved in DMSO (Sigma-Aldrich) or 200nM of the membrane impermeable fluorescent dye SYTOX Green dissolved in H_2_O (Thermo Fisher) for 5 min prior to imaging. 1% DMSO was maintained upon staining with DiSC_3_(5) to aid dye solubility. For the determination of the membrane potential using DiSC_3_(5), the cells were diluted to an OD_600_ of 0.3 (the lowest density tested) to ensure a similar cell:dye ratio. The cells were diluted using spent medium obtained by centrifugation of an aliquot from the same culture. As positive controls for membrane depolarisation or membrane permeabilisation, the cells were incubated with either 10μM of gramicidin or 10μM nisin for 5 min prior to imaging . Incubations with dyes or antibiotics were performed by transferring 200 μl of cells into a 2 ml microcentrifuge tube with a perforated lid (to allow aeration), and shaking at 850 rpm and 37°C in a ThermoMixer (Eppendorf) for 5 min. For visualisation, the cell suspensions were immobilised on Teflon-coated multi-spot microscope slides (Hendley-Essex) covered by a thin layer of 1.2% (w/v) agarose. For membrane potential measurements, the slide was pre-warmed to 37°C prior to imaging. A sample of the cell suspension (0.5 μl) was applied to the agarose surface, air-dried, and covered with a coverslip. The slides prepared in this manner were imaged within 5 min of the application of the coverslip to minimise oxygen limitation and the associated loss of membrane potential. See te Winkel et al. (23) for further details regarding slide preparation and time-lapse microscopy. The phase contrast and fluorescence imaging were carried out using a Nikon Eclipse Ti microscope (Nikon Plan Apo 100x/1.40 Oil Ph3 objective) equipped with Semrock Cy5-4040C (EX 628/40, DM660lp, EM 692/40) and Chroma 49002 (EX470/40, DM495lpxr, EM525/50) filter sets for imaging DiSC_3_(5) and SYTOX Green, respectively. The images were acquired using Metamorph 7.7 (Molecular Devices, inc).

### Microscopy analysis

For fluorescent microscopy, images were analysed using the Fiji software package (29). Quantification of DiSC_3_(5) fluorescence was performed in a semi-automated manner . In brief, the individual cells were identified as regions of interest (ROi) by thresholding the phase-contrast images . If cells grew as clusters that could be visually identified as individual cells but that could not be separated based on image thresholding, the ROis were separated by a manually drawn thin line. The respective fluorescence images were background-subtracted prior to analysis. The mean fluorescence intensity values for single cells were acquired using a custom script available at https://github.com/NCL-imageAnalysis/General_Fiji_Macros.

### Fluorometric measurement of lysis using SYTOX Green

For the determination and quantification of cell lysis in a *B. subtilis* culture during growth for 15 h, culture supernatant containing the lysis-derived DNA was collected by centrifuging 500 μl of culture in a 1.5 ml Eppendorf tube (5 min, RT, 16,000 x*g*). The supernatant was diluted 1:10 in a black, polystyrene, flat-bottomed microtiter plate (Porvair Sciences), and 1μM of the DNA-intercalating dye SYTOX Green (Thermo Fisher Scientific) was added and incubated while shaking at 200 rpm at room temperature in a BMG Clariostar multimode plate reader using 485-10 nm excitation and 520-10 nm emission wavelength windows, respectively. Fluorescence intensity was monitored until stable levels were obtained. The background fluorescence of SYTOX Green in medium-only samples was subtracted from the test samples. For the preparation of the SYTOX Green-based lysis measurements, a calibration curve was generated from samples of *B. subtilis* grown as described above to different cell densities. Culture samples (500 μl) were collected at various optical densities and sonicated on ice for 10 min to lyse the cells and release cellular DNA. The supernatants of the sonicated samples were subjected to the fluorometric SYTOX Green assay as described above. The resulting calibration curve was used to convert the measured SYTOX Green signal to the pre-lysis OD_600_ of the cell population, assuming full release of the cellular DNA content into the culture supernatant. The percentage of cell lysis was then calculated as follows: Percentage cell lysis = (calculated pre-lysis OD_600_ of the lysing cell population)/(OD_600_ of culture)*100.

### Statistical analysis

Statistical analysis was performed using GraphPad Prism 9.5, making use of either an ordinary one-way, unpaired ANOVA with a Dunnett’s multiple comparisons test or an unpaired, two-sided t-test.

## Results

### Use of a membrane permeability indicator to monitor autolysis in a *B. subtilis* culture

If the majority of cells in a non-growing bacterial culture undergo lysis, this can simply be monitored by the decline in culture optical density. However, if the lysis process only affects a subpopulation of a growing culture, determining the extent of lysis is less trivial and the associated reduced rate of optical density increase can easily be misinterpreted as a reduction in the growth rate of intact cells. An alternative to monitoring changes in culture optical density is the use of so-called viability indicators or Live/Dead stains such as SYTOX Green or propidium iodide, which detect cells with compromised membrane integrity (19, 20). These assays rely on DNA-intercalating dyes that exhibit strongly enhanced fluorescence upon DNA binding but are membrane impermeable and thus unable to stain the DNA of cells with intact cell membranes. While these dyes are more commonly used to detect the activity of membrane-disrupting compounds (25, 30), the integrity of a bacterial cytoplasmic membrane is also disrupted as part of the autolytic process, thus potentially allowing detection of autolysis as well. However, to facilitate a robust interpretation of lysis data, it is crucial to consider the fate of cytoplasmic DNA during the autolytic process.

CCCP is an ionophore that transports protons across the bacterial cytoplasmic membrane, thus triggering a collapse of the proton motive force (pfm) and an associated decline in cell ATP levels in *B. subtilis* (10). This comprehensive de-energisation induces misregulation of cell wall hydrolases, ultimately leading to induced autolysis (31, 32). To monitor the fate of cellular DNA following *B. subtilis* autolysis, we followed CCCP-induced lysis using time-lapse microscopy and SYTOX Green staining. As shown in Figure 1a and Movie S1, autolysis indeed triggers permeabilisation of the cytoplasmic membrane that can be detected with SYTOX Green prior to significant changes in phase contrast becoming evident. However, the fate of the cellular DNA is rather heterogeneous with both rapid release to the medium and retention within the lysed cell remnants being observed. Similar heterogeneity is evident when autolysis is induced with the pore-forming antimicrobial peptide Nisin (Figure 1b), which forms large pores in *B. subtilis* membranes in a lipid ii-dependent manner (33). While the pore-forming activity allows SYTOX Green to rapidly enter the cell and stain cytoplasmic DNA, prolonged incubation leading to autolysis is associated with at least partial release of cellular DNA . In conclusion, while SYTOX Green fluorescence can be used to monitor the autolytic process, the associated release of cellular DNA complicates single-cell analysis unless time-lapse microscopy is used. Nonetheless, SYTOX Green does gain access to DNA upon autolysis, which should enable the development of a population-based bulk assay.

### Analysing autolysis with a membrane permeability indicator in culture-based bulk assays

SYTOX Green and Propidium iodide (Pi) are frequently used in fluorescence plate reader-based bulk assays to measure changes in membrane permeability as a proxy for cell viability (19, 20, 33). However, these assays are typically used to monitor a relatively rapid response to compounds and conditions that directly affect membrane-permeabilising properties. When we monitored SYTOX Green and Pi fluorescence directly in *B. subtilis* cultures grown into the late post-exponential phase using a fluorescence plate reader, no clear signature for autolysis was detected even though we expected the post-exponential, nutrient-limited cultures to exhibit significant levels of autolysis (Supplementary Figure 2). Thus, autolysis affecting part of the *B. subtilis* cell population cannot be easily detected by monitoring changes in the SYTOX Green and Pi fluorescence of the culture.

To confirm that late post-exponential phase *B. subtilis* cultures do indeed undergo partial autolysis, we determined whether DNA released upon lysis could be detected in culture supernatants (Figure 2a). This was achieved by staining a culture sample with SYTOX Green and assaying the SYTOX Green fluorescence signal. While there was a strong fluorescence signal in the culture supernatant fraction, only a weak signal was detected in the pellet fraction (Figure 2b), presumably due to the release of the cellular DNA linked to the forces involved in compacting intact cells to high cell density (34). This suggests that a significant degree of autolysis is indeed occurring in such a *B. subtilis* culture, and that most of the DNA of lysed cells ultimately accumulates in the culture supernatant rather than remaining trapped within lysed cells. Finally, the presence of DNA in the culture supernatant was further confirmed by running the sample on an agarose gel stained with the DNA dye Nancy-520 (Figure 2c).

**Figure 2:**
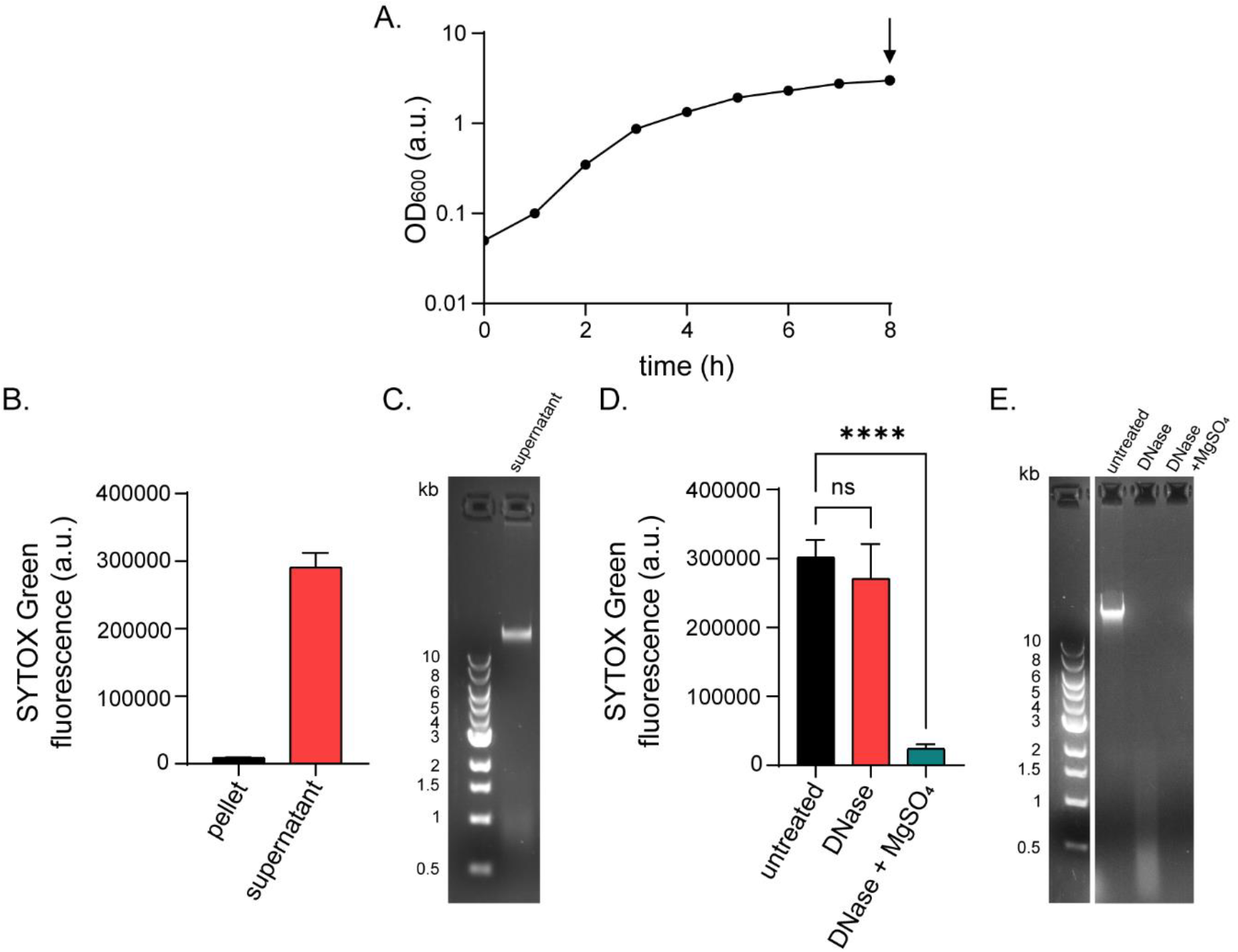
Extracellular DNA is present in the growth medium of post-exponential growth phase *B. subtilis* cultures. **(A)** Growth kinetics (OD_600nm_) of *B. subtilis* grown in mSMM at 37°C with the black arrow indicating the time point of sample collection. **(B)** SYTOX Green fluorescence signal measured fluorometrically for the cell pellet and the corresponding culture supernatant. The graph depicts the mean and standard deviation from three independent biological replicates. **(C)** Agarose gel depicting DNA stained with Nancy-520 in one of the supernatant samples analysed in panel B. **(D)** The SYTOX Green fluorescence signal for an undigested culture supernatant sample, and for samples digested with 10 Kunitz units of bovine pancreas DNase i for 1 h at 37°C in the presence and absence of a supplemented divalent cation (20 mM MgSO4). The graph depicts the mean and standard deviation from three independent biological replicates, along with P values of a one-way, unpaired ANOVA. **(E)** Agarose gel depicting DNA stained with Nancy-520 in the same supernatant samples as shown in panel D. The 1 kb DNA Ladder from NEB^®^ was used as a molecular size marker. Strain used: *B. subtilis* prototroph (MDS23). **** = p ≤ 0.0001; ns = no significant difference.

To verify that the SYTOX Green signal detected in the culture supernatant was due to the presence of DNA, the supernatant samples were incubated with endoribonuclease DNase I in the presence or absence of supplementary Mg^2+^, an essential cofactor for activity (35, 36). When supernatant samples were incubated with DNase I in the absence of supplementary Mg^2+^, a fainter smear of lower molecular sized DNA was observed. However, no significant reduction of the corresponding SYTOX Green signal was observed (Figure 2d, e) . In contrast, when supplementary Mg^2+^, was added to the supernatant along with DNase i, the SYTOX Green signal was significantly reduced and no DNA was detected in the agarose gel, indicating extensive DNA degradation (Figure 2d, e). Thus, the SYTOX Green signal detected in culture supernatant is specific for DNA.

Endogenous nucleases can potentially degrade DNA released when cells lyse. *B. subtilis* encodes two major and one minor extracellular nuclease: the membrane-associated nuclease NucA, involved in DNA cleavage during transformation (37, 38), the sporulation-specific secreted nuclease NucB (39, 40) and the minor biofilm-associated YhcR (41). When SYTOX Green culture supernatant signals were measured for strains lacking the NucA and NucB extracellular nucleases, no significant differences in extracellular DNA levels were measured compared to the wild-type culture (Supplementary Figure 3). Thus, the presence of the two major *B. subtilis* extracellular nucleases had no significant, measurable effect on the observed level of DNA in the culture supernatant throughout the growth phases. YhcR was not tested as it is only induced in response to the need to recover eDNA from biofilm pellicles (41). Moreover, when purified *B. subtilis* genomic DNA was artificially fragmented by sonication, it had little effect on the measured SYTOX Green fluorescence signal (Supplementary Figure 4). Together, these results confirm that the SYTOX Green signal in the culture supernatant provides a sensitive assay for cell lysis that is largely insensitive to DNA fragment size.

Finally, to verify that the observed extracellular DNA is indeed autolysis-derived, early post-exponential growth phase culture samples from a strain deficient for several autolytic enzymes (*B. subtilis* Δ*lytABCDEF*), and from its immediate parent strain (*B. subtilis* 168) were collected and analysed (Figure 3a). Very little DNA was observed in the culture supernatant of the autolytic mutant based on SYTOX Green fluorescence (Figure 3b), and no DNA was detectable in the corresponding agarose gel (Figure 3c). These results confirm that the DNA present in the *B. subtilis* culture supernatant is indeed derived from cells undergoing autolysis in a culture that, at population level, exhibits healthy and robust growth.

**Figure 3:**
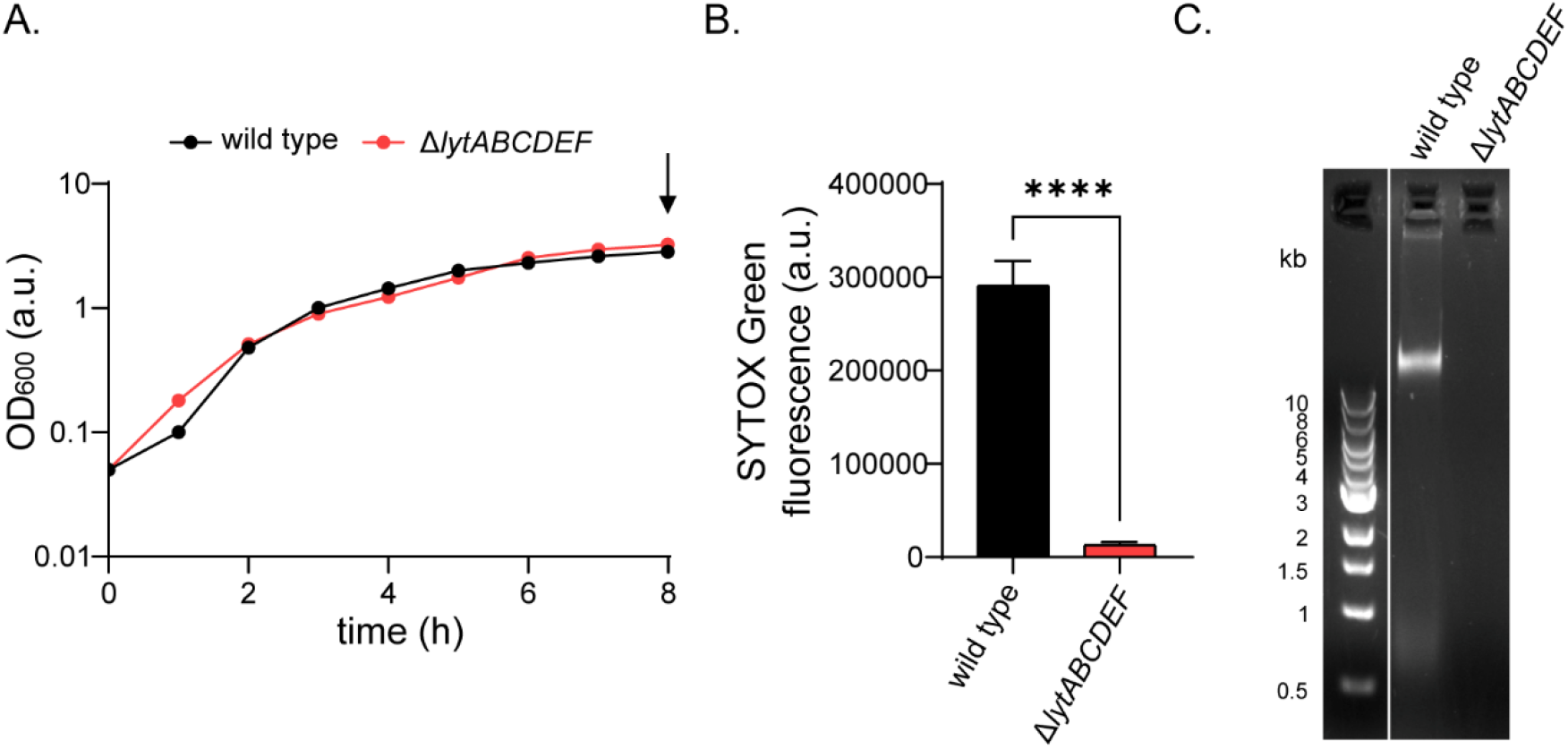
Extracellular DNA present in *B. subtilis* culture supernatants is lysis-derived. **(A)** Growth kinetics (OD_600nm_) of *B. subtilis* wild type (168) and the filamentous multiple autolytic enzyme-deficient derivative (Δ*lytABCDEF*) strain in mSMM at 37°C. The black arrow indicates the time point of sample collection. **(B)** SYTOX Green fluorescence signals of corresponding culture supernatant samples. The graph depicts the mean and standard deviation from three independent biological replicates. **** = p ≤ 0.0001. **(C)** Agarose gel depicting DNA stained with Nancy-520 of one of the supernatant samples indicated in panel A and measured in panel b The 1 kb DNA Ladder from NEB^®^ was used as a molecular size marker. Strains used: *B. subtilis* 168 (wild type) and *B. subtilis* KS19 (Δ*lytABCDEF*).

### Quantification of lysis in a growing *B. subtilis* culture

As shown above, SYTOX Green fluorescence is insensitive to the extent of fragmentation of lysis-derived DNA. This observation, combined with the lack of significant degradation by the two major extracellular nucleases, means that the accumulation of extracellular DNA should serve as a cumulative measure of the lysis occurring in a culture. To estimate the extent of cell lysis, the SYTOX Green fluorescence signal was calibrated using cell suspensions of known optical density that were sonicated to release their cytoplasmic DNA into the supernatant. The respective samples were then stained with SYTOX Green, quantified using a fluorescence plate reader, and the SYTOX Green fluorescence intensities were plotted against the sample OD_600_ values (Figure 4a). A linear correlation between the measured SYTOX Green fluorescence and corresponding OD_600_ values (0-1.2) was observed (Figure 4a, b). This calibration facilitates a simple conversion between the measured supernatant SYTOX Green fluorescence intensity and the cumulative fraction of the cell population that has released the observed extracellular DNA through lysis.

**Figure 4:**
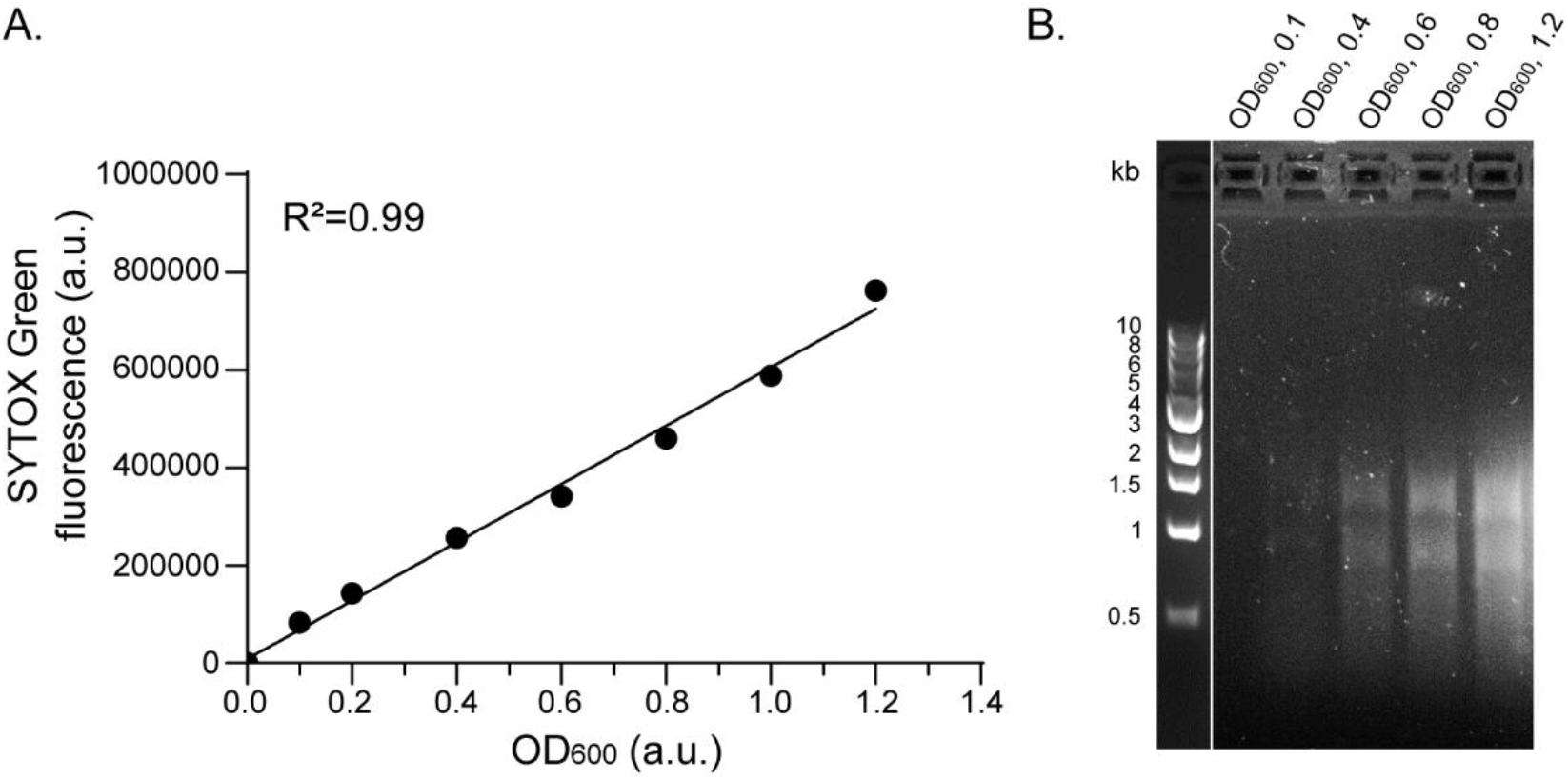
Calibration of the SYTOX Green autolysis assay. **(A)** SYTOX Green fluorescence signals of a *B. subtilis* culture grown to various optical densities in mSMM and subsequently lysed by sonication to release DNA, plotted against the OD600 of the samples prior to sonication. **(B)** Agarose gel showing the supernatant DNA stained with Nancy-520 in the same samples shown in panel A. The 1 kb DNA Ladder from NEB^®^ was used as a molecular size marker. Strain used: *B. subtilis* prototroph (MDS23).

### Establishing the DiSC_3_(5) assay for measuring single-cell energy levels from *B. subtilis* cultures of varying growth phases and cell densities

The membrane potential-sensitive carbocyanine dye 3,3′-dipropylthiadicarbocyanine iodide, DiSC_3_(5), is a well-established reporter for measuring bacterial membrane potential and, therefore, a sensitive single-cell reporter for the general bioenergetic status of various bacterial species, including *B. subtilis* (23, 25, 42). However, its use has been primarily limited to low-density, exponential growth phase cultures, and to comparing cultures with identical optical densities. The membrane potential-sensitive staining by DiSC_3_(5) is due to its hydrophobic character, which allows it to diffuse across cell membranes, and the cationic charge, which enables transmembrane electric fields to bias its diffusion. As a result, DiSC_3_(5) acts as a so-called Nernstian dye, establishing a gradient across the bacterial cytoplasmic membrane that scales with the membrane potential (24, 43, 44). Due to this mechanism of membrane potential sensing, the single-cell DiSC_3_(5) fluorescence levels are influenced by cell sensitivity and experimental factors that can rapidly alter the cellular metabolic state, such as changes in nutrient supply. A common experimental pitfall in this context is the washing and resuspension of cells in a nutrient-free buffer, which results in rapid loss of membrane potential (40).

To establish a robust assay for measuring single-cell membrane potential levels for varying growth phases and cell densities, the impact of cell density was first analysed. Accordingly, *B. subtilis* was grown to an OD_600_ of 0.6 (exponential phase) and then diluted to an OD_600_ of 0.45, 0.3 or 0.15. To prevent re-energisation of cells upon resuspension in fresh medium, cells were diluted in culture supernatant obtained by centrifugation of a parallel sample from the same culture.

These results demonstrate that the single-cell DiSC_3_(5) levels are highly sensitive towards sample optical density, with diluted samples exhibiting stronger and more homogeneous staining levels despite originating from the same culture (Figure 5a, b). Thus, to reliably measure and compare membrane potential in *B. subtilis* cultures of varying cell densities, it is necessary to equalise the sample optical density . In our experimental conditions, diluting to an OD_600_ of 0.3 is sufficient to ensure homogeneous DiSC_3_(5) staining levels, although these conditions should be verified when applied to different bacterial species and media. Using cell-free “spent” media obtained from corresponding cultures is strongly recommended to minimise changes in nutrient availability.

**Figure 5:**
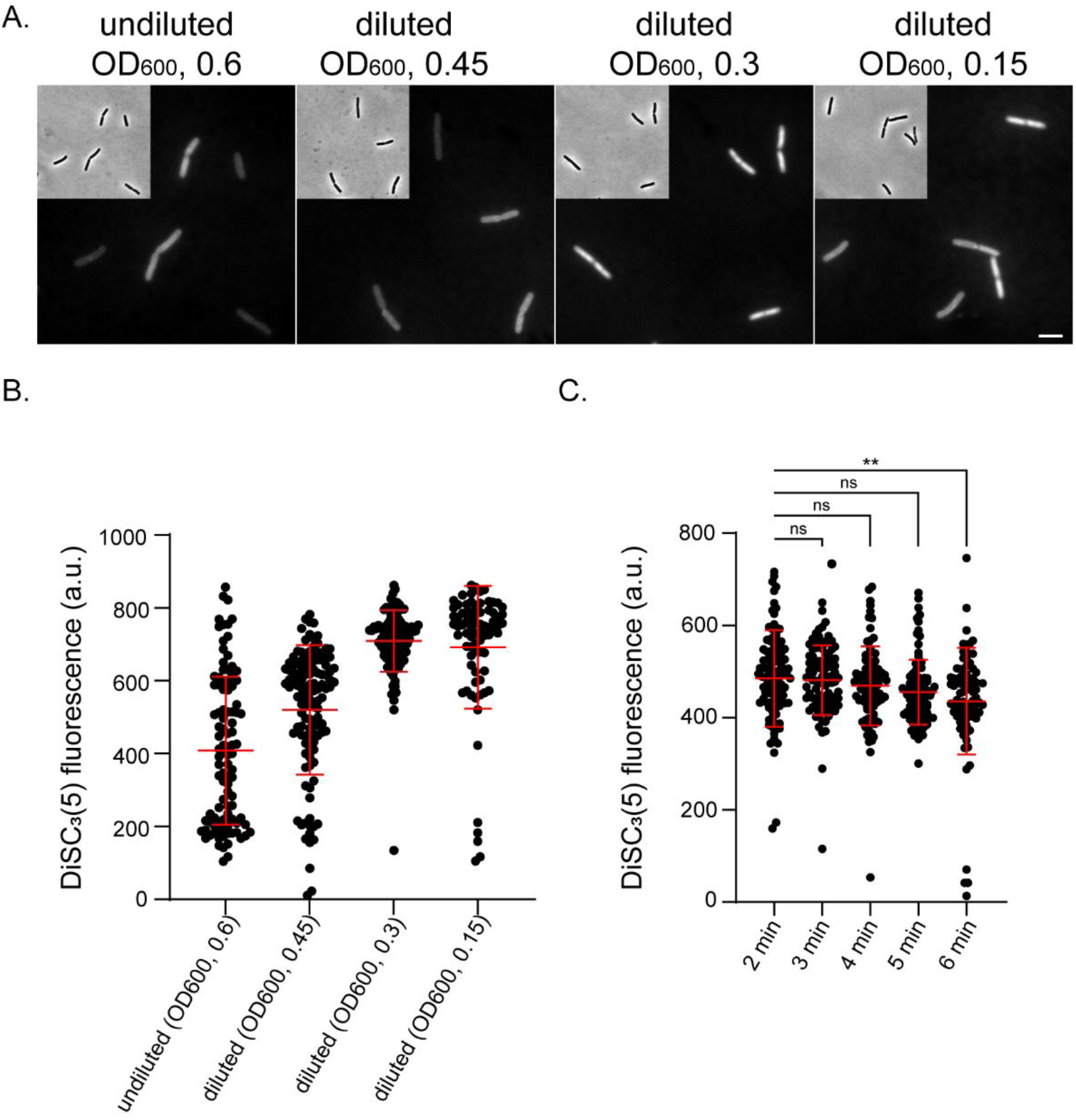
Cell densities and imaging speed influence *B. subtilis* DiSC_3_(5) fluorescence signals. **(A)** Phase contrast and fluorescence microscopy images of DiSC_3_(5)-stained *B. subtilis* in mSMM in undiluted (OD_600_ of 0.6) and diluted cultures (OD_600_ of 0.45 to 0.15). **(B)** Quantification of DiSC_3_(5) fluorescence intensities for individual cells from the imaging dataset shown in panel (a) (n=100). **(c)** Quantification of DiSC_3_(5) fluorescence intensities for individual cells (n=90), 2-6 min after immobilisation of cells on microscope slides (application of the coverslip). Mean fluorescence intensities and standard deviation are indicated with red lines, together with P values of a one-way, unpaired ANOVA. ** = p ≤ 0.01; ns = a non-significant difference. Scale bar: 3 μm. Strain used: *B. subtilis* prototroph (MDS23).

It has previously been shown that *B. subtilis* cells are affected by oxygen limitation under microscope cover slips and, as a result, gradually lose membrane potential when immobilised on a microscopy slide (10, 11). To establish a timeframe in which DiSC_3_(5) microscopy can be applied for assessing single-cell energy levels, a *B. subtilis* culture was grown to an OD_600_ of 0.3, stained with DiSC_3_(5), placed on the agar-coated microscope slide, covered with a cover slip, and imaged every minute up to 6 min (Figure 5c). At this sample cell density, *B. subtilis* can maintain stable membrane potential levels under the cover slip for approximately 5 min. After this time window, the membrane potential starts to decline and become more heterogeneous . In conclusion, it is possible to measure membrane potential levels reliably for relatively dilute cell suspensions (OD_600_ = 0.3), provided the data are acquired within 5 minutes of immobilisation.

### *B. subtilis* wild type undergoes autolysis and exhibits strong energy level heterogeneity in glucose-based minimal medium

After establishing the DiSC_3_(5) assay for measuring single-cell energy levels from *B. subtilis* cultures of varying growth phases and cell densities, and the SYTOX Green assay for quantifying cumulative lysis by measuring extracellular DNA, we investigated individual cell membrane potential levels and cell lysis throughout the growth cycle. As shown previously (Figure 1a), the de-energisation of *B. subtilis* by collapsing the proton motive force with CCCP induces cell lysis. While the exact molecular mechanism that links cell de-energisation to cell lysis is poorly understood, it is well established that the process involves misregulation of the cell’s own autolytic enzymes (9, 43). Misregulation is induced by numerous compounds that compromise *B. subtilis* membrane integrity or hamper the cell’s ability to maintain its proton motive force by respiration (45). To gain greater insight into the extensive accumulation of lysis-derived extracellular DNA in an apparently healthy, well-growing *B. subtilis* culture, we monitored and correlated the extent of single-cell membrane potential heterogeneity and cell lysis throughout the growth cycle of *B. subtilis*.

When *B. subtilis* was grown in a glucose-based minimal medium (mSMM), the level of DNA that accumulated in the supernatant during and towards the end of exponential growth was ∼5% of the DNA associated with the cells (% Lysis; Figure 6a, T_0_) . In contrast, upon slow transition from the exponential to the post-exponential growth phase (so-called transition phase T_0_), the cumulative cell lysis increased to ∼19% by T_+4_ despite the continuous increase in culture optical density. This high level of background lysis was further confirmed by agarose gel electrophoresis whereby increasing amounts of extracellular DNA were detected over time (Figure 6b). When the corresponding single-cell energisation levels were monitored using membrane potential as a proxy, *B. subtilis* was found to be well energised throughout the exponential growth phase, with little single-cell heterogeneity observed (Figure 6c, T_-4_ to T_-1_). However, as the growth rate declined and the culture transitioned to the post-exponential growth phase, the average single-cell membrane potential levels gradually declined (Figure 6c, T_0_ to T_+4_). Strikingly, the cell population now began to exhibit high energy level heterogeneity with the majority of intact (phase dark) cells maintaining reasonably well-energised membranes while a sub-population of such cells (∼3-6%) demonstrating near or complete de-energisation. The largest number of de-energised cells was present at T_+4_ h, with ∼31% of cells exhibiting very low membrane potential. Hence, there is a striking correlation between the emergence of a de-energised subpopulation and the observed increase in autolysis. It is tempting to speculate that the observed lysis is indeed due to the emergence of a de-energised cell population that ultimately undergoes autolysis. This raises the question of whether this is programmed altruistic response, designed to provide nutrients to help sustain the population of energised cells, or a stochastic response to decreasing nutrient availability. More detailed studies are needed, however, to confirm this link, to decipher the physiological processes responsible for the high level of membrane potential heterogeneity observed in post-exponential *B. subtilis* cultures and to devise strategies to maintain productivity in high-density industrial fed-batch cultures.

**Figure 6:**
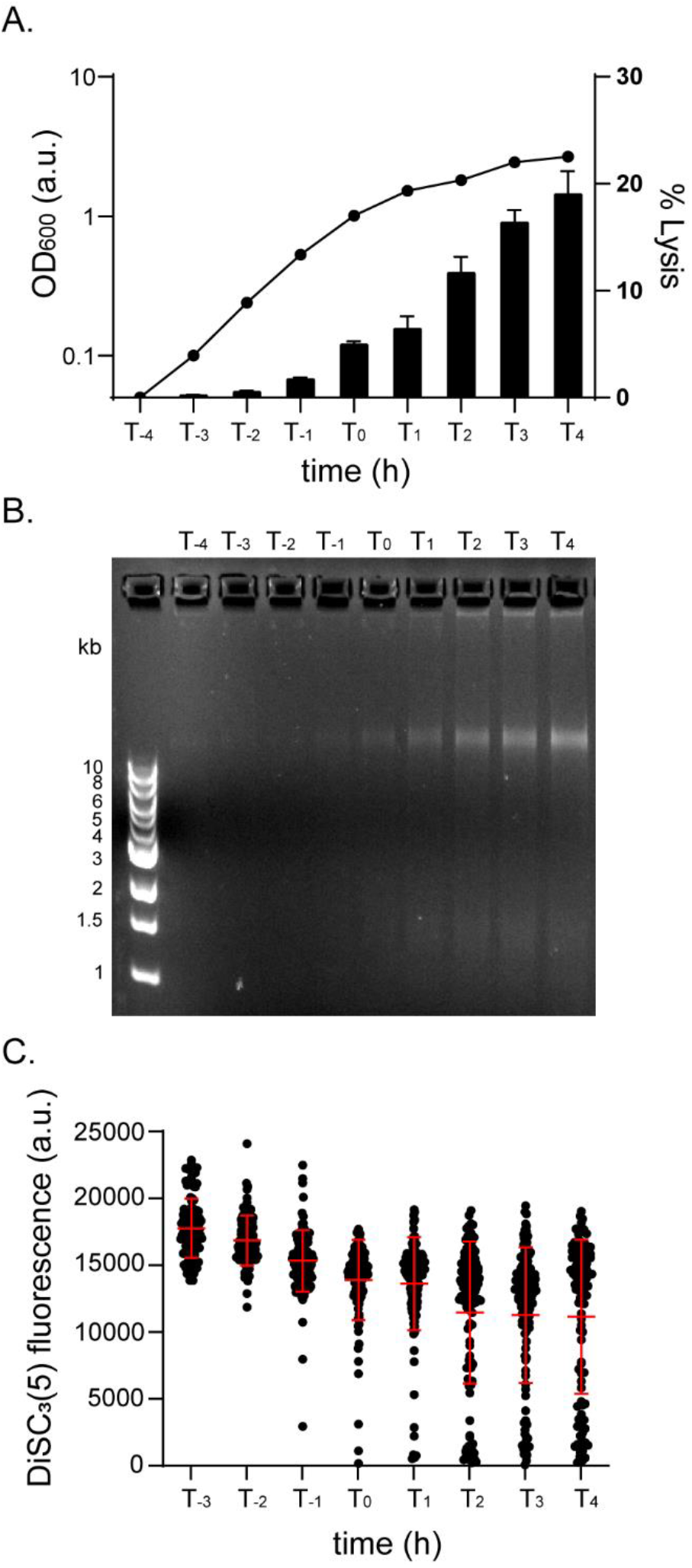
*B. subtilis* exhibits growth phase-dependent autolysis and energy level heterogeneity. **(A)** Growth kinetics (OD_600nm_) of *B. subtilis* in mSMM along with the percentage of cells cumulatively lysed at specified timepoints, calculated using the calibration curve shown in Fig. 4. **(B)** Agarose gel depicting DNA stained with Nancy-520 in the same lysis-derived DNA samples shown in panel A. The 1 kb DNA Ladder from NEB^®^ was used as a molecular size marker. **(C)** Quantification of DiSC_3_(5) fluorescence intensities for individual cells (n=103-128) at the specific time points before and after transition (T0) from exponential towards post-exponential phase growth. Mean fluorescence intensities and standard deviations are indicated with red lines. Strain used: *B. subtilis* prototroph (MDS23).

## Discussion

The production of biologicals by microbial fermentation broadly falls into two categories . In the first case the desired product accumulates in the cytoplasm of the production strain and therefore must be released by controlled cell lysis at the end of fermentation. This means of production primarily suits higher-value products, since the required downstream purification costs are generally high. The alternative involves secreting products such as proteins, enzymes and metabolites from intact cells into the culture from where they are extracted more cost-effectively following cell removal . In the latter case a significant level of cellular lysis during the production phase is a major issue affecting downstream processing costs, since it releases cell-associated proteases that reduce product yield and undesirable cellular components, such as endotoxins, lipids and nucleic acids that must be removed prior to regulatory approval.

Developing genetic and processing strategies to reduce lysis requires assays that allow the metabolic status of the cells and the extent of lysis to be monitored efficiently. In this manuscript, we provide experimental guidance and protocols to extend the usability of two physiological fluorescence reporters, the voltage-sensitive dye DiSC_3_(5) and the membrane permeability indicator SYTOX Green, in the Gram-positive industrial bacterium *B. subtilis*. With the approaches detailed here, DiSC_3_(5) can be used to monitor bacterial single-cell energisation levels robustly across varying growth phases and cell densities, thereby significantly expanding its usefulness for bacterial physiological and bioenergetic studies. Furthermore, we have established how the membrane permeability indicator SYTOX Green can be used as a reporter for autolysis at the single-cell level and to quantitatively monitor cell lysis affecting a subpopulation of cells in an otherwise growing culture. While the assays are established for *B. subtilis*, both DiSC_3_(5) and SYTOX Green can be used in a variety of bacterial species (20–23, 42, 46). As a result, establishing a similar assay for other bacterial species should be relatively straightforward, building on the protocols provided here for *B. subtilis*.

Gross measurements of culture density increases during fermentation do not reflect activity at the single-cell level, but rather the balance between cell mass increase, cell death and lysis. Once we had established the detailed assays for *B. subtilis* cultures, we encountered increasingly high levels of membrane potential heterogeneity during transition to and in the post-exponential growth phase. To our knowledge, this degree of heterogeneity in energisation levels has not been previously observed. One cellular consequence of this population heterogeneity could indeed be the unexpected levels of cell lysis observed in dense *B. subtilis* cultures. Under these conditions, *B. subtilis* undergoes differentiation into various subpopulations with distinct physiological characteristics and roles (47). The observed heterogeneity in energisation and cell lysis may be link to the developmental programmes that underpin *B. subtilis* differentiation, providing a source of nutrients when exogenously supplied nutrients are limited. While we are currently actively studying the underlying molecular mechanism and its consequences for *B. subtilis* physiology, details regarding this underappreciated biological phenomenon are beyond the scope of this manuscript.

## Acknowledgements

We thank James Grimshaw for his support with image analysis.

## Author contributions

Authors’ contributions: M.D.S and M.H. performed the experiments, M.D.S, T.v.R, C.R.H and H.S designed experiments and helped with data interpretation, T.v.R, C.R.H and H.S conceived and supervised the project, M.D.S, C.R.H and H.S wrote the manuscript.

## Availability of data and materials

Source data for all figures and graphs presented in the manuscript are available via Newcastle University’s research data repository via https://doi.org/10.25405/data.ncl.29924237. The strains and plasmids are available upon request to H.S

## Declarations

## Ethics approval and consent to participate

Not applicable

## Consent for publication

All authors have consented to the publication of this manuscript.

