## Supplemental material for "Monitoring the bioenergetic state and cell lysis of *Bacillus subtilis* throughout the growth cycle"

**Monitoring single cell bioenergetic status and cell lysis in dense and differentiating *Bacillus subtilis* cultures**

Supplementary Figures 1-4

Supplementary Movie S1

**SUPPLEMENTAL MATERIAL**

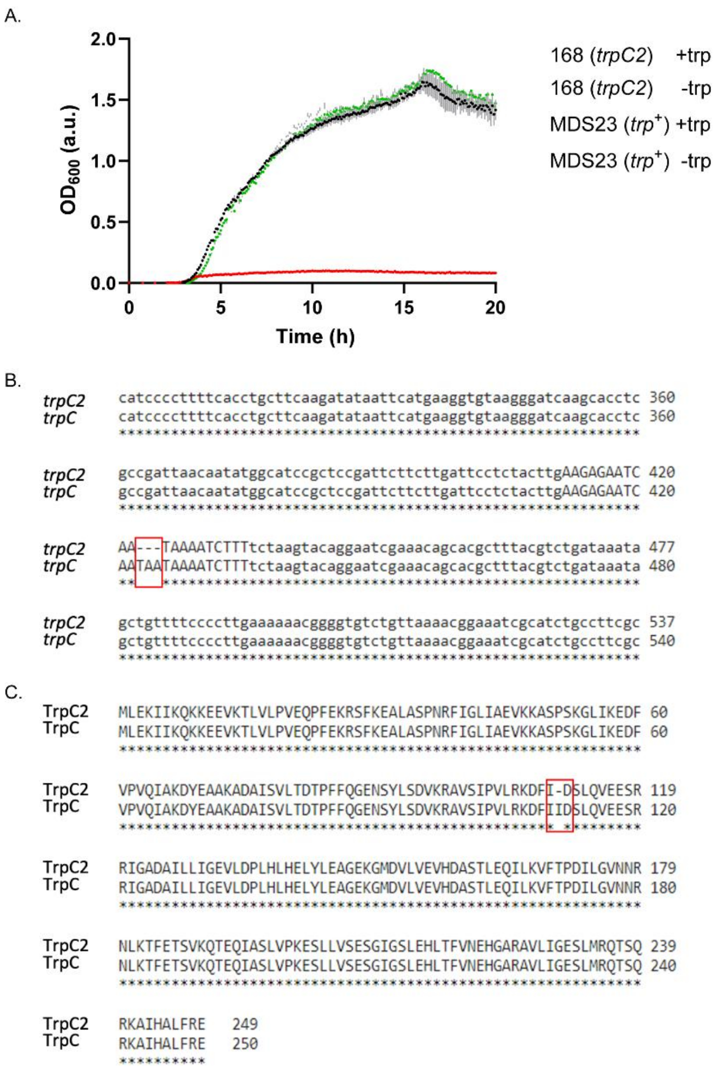


**Supplementary Figure 1: *B. subtilis* strain MDS23 was generated by transforming strain 168 (*trpC2*) with a PCR-generated DNA from the *trpC* gene of NCIB3610$^{T}$ and selection for prototrophic transformants.**

**(a)** Growth of strains 168 and MDS23 in modified SMM + glucose (0.5% v/v) with and without tryptophan (0.002% v/v). The growth of MDS23 without tryptophan is identical to that of 168 with tryptophan, confirming the loss of tryptophan auxotrophy. The growth data is the average of three independent cultures. **(b)** The nucleotide sequence of the *trpC2* gene in strain 168 compared to the prototrophic (*trpC*) version of the gene in MDS23. The red box marks the missing TAA codon. **(c)** The derived amino acid sequences of the TrpC2 and TrpC proteins of strains 168 and MDS23, respectively. The sequence shows the position of the missing Ile (red box) at position 111, located at the junction of the β3 strand and α3 helix. Although not directly involved in the formation of the active site, the deletion abolishing the activity of the enzyme (50).

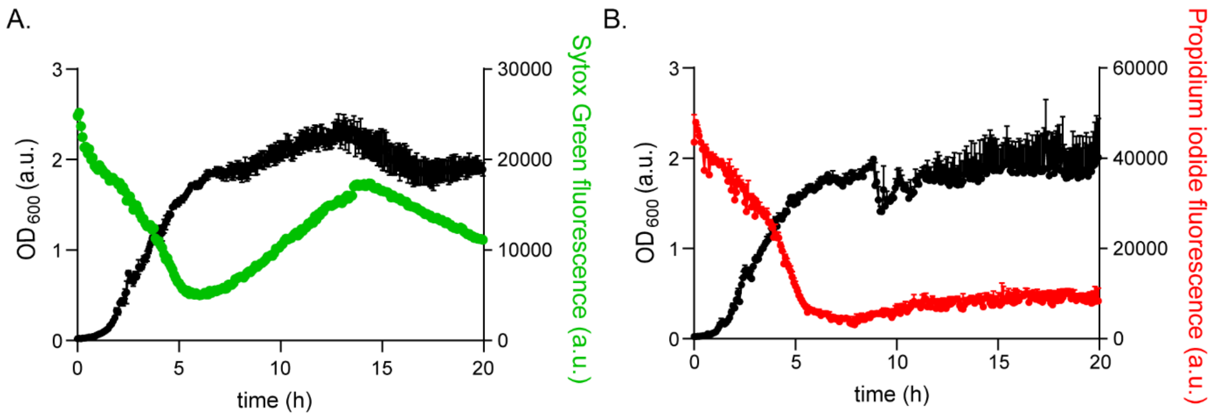


**Supplementary Figure 2: Continuous monitoring of lysis in a growing *B. subtilis* culture using SYTOX Green or Propidium Iodide is not feasible.**

Growth and fluorescence measurements of *B. subtilis* in mSMM with the presence of either **(a)** 1 µM SYTOX Green or **(b)** 10 µM propidium iodide. Graphs depict the means and standard deviation from three technical replicates. Strain used: *B. subtilis* prototroph (MDS23).

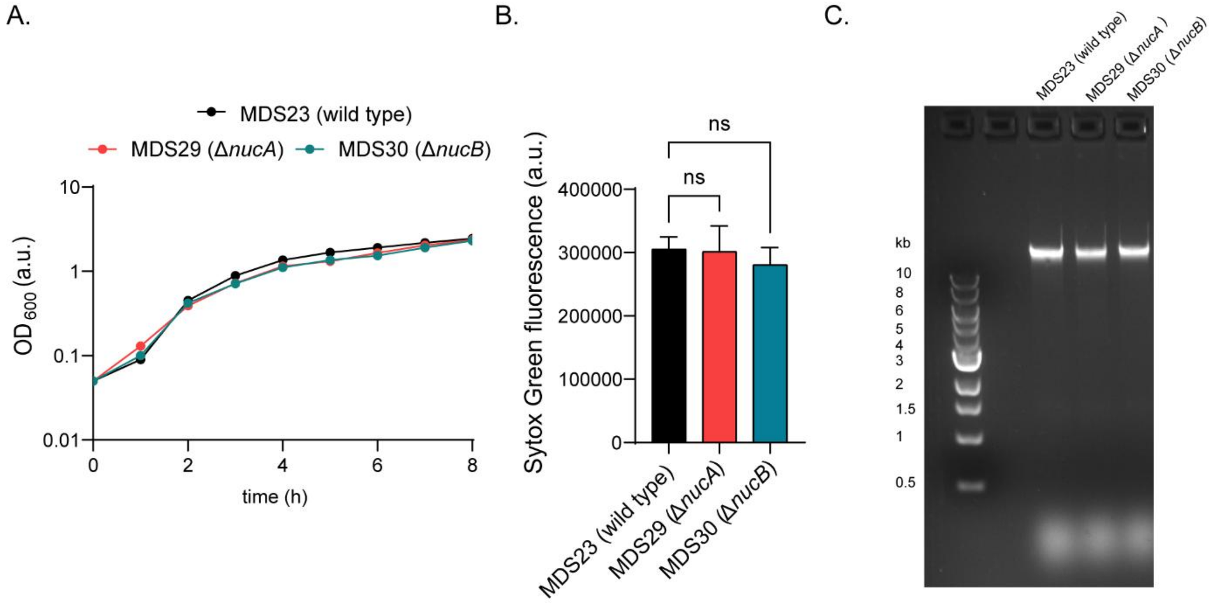


**Supplementary Figure 3: The deletion of two major *B. subtilis* nucleases does not significantly affect culture supernatant DNA levels**

**(a)** Growth kinetics of wild type, Δ*nucA* and Δ*nucB* in mSMM at 37°C with the black arrow indicating the time point of sample collection. **(b)** SYTOX Green fluorescence signal of supernatants derived from an early stationary phase wild type, Δ*nucA* and Δ*nucB* cultures. The graph depicts mean and standard deviation from three independent biological replicates, together with P values of a one-way, unpaired ANOVA. ns indicates a non-significant difference. **(c)** Agarose gel depicting DNA stained with Nancy-520 one of the supernatant samples shown in panel A. The 1 kb DNA Ladder from NEB® was used as a molecular size marker. Strains used: *B. subtilis* prototroph (MDS23), Δ*nucA* (MDS29) and Δ*nucB* (MDS30).

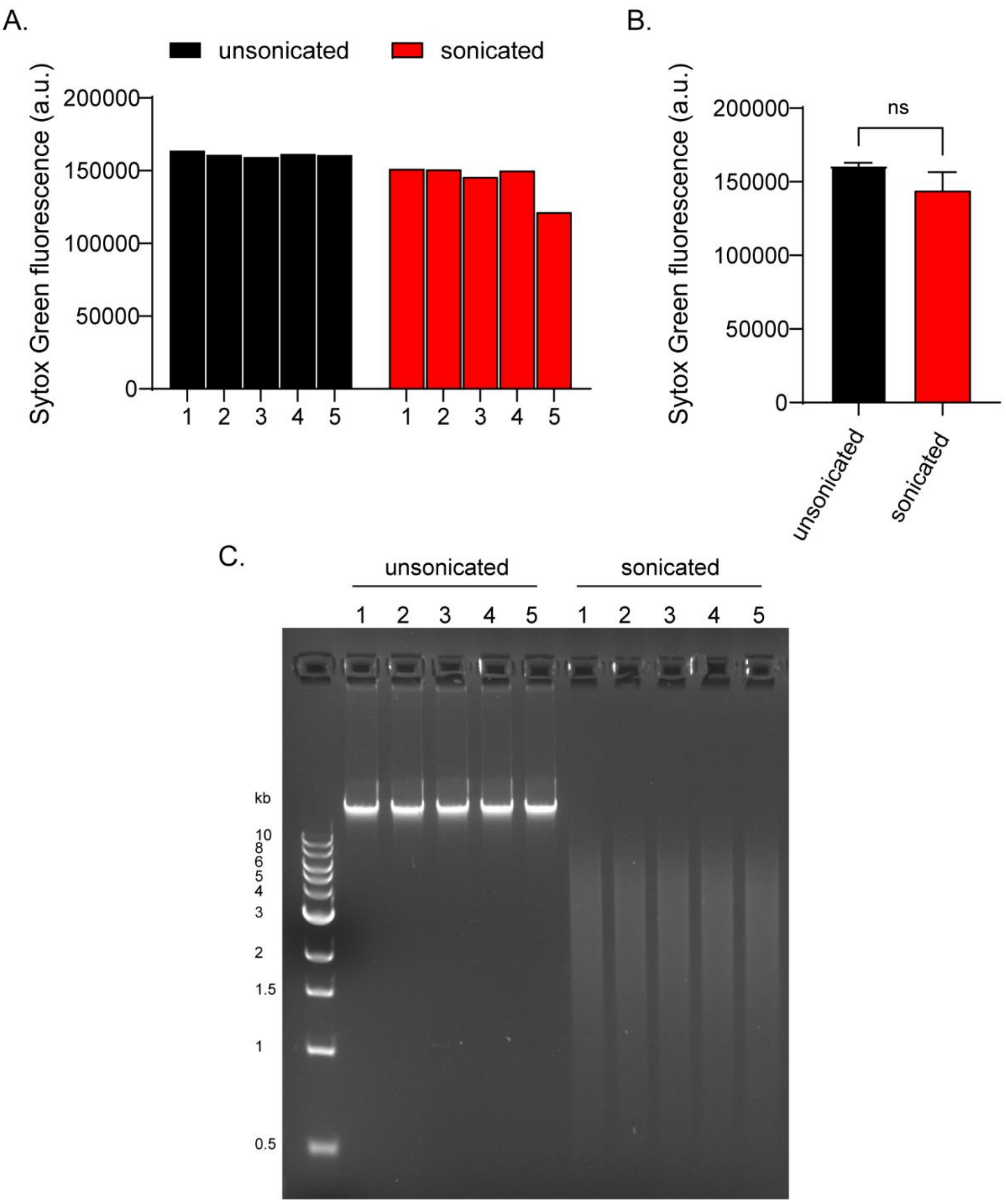


**Supplementary Figure 4: Fragmentation of DNA by sonication does not significantly affect the SYTOX Green fluorescence signals**

**(a)** SYTOX Green fluorescence signal of 5 technical replicates of *B. subtilis* wild type genomic DNA before and after sonication at 40% amplitude for 10 min. **(b)** The graph depicts the mean and standard deviation of 5 technical replicates, along with the P values of a one-way, unpaired ANOVA. ns indicates a non-significant difference. **(c)** Agarose gel depicting DNA stained with Nancy-520 in the samples shown in panel **(a)**. The 1 kb DNA Ladder from NEB® was used as a molecular size marker. The contrast setting of the agarose gel image has been altered to enhance the visualisation of heavily fragmented DNA.

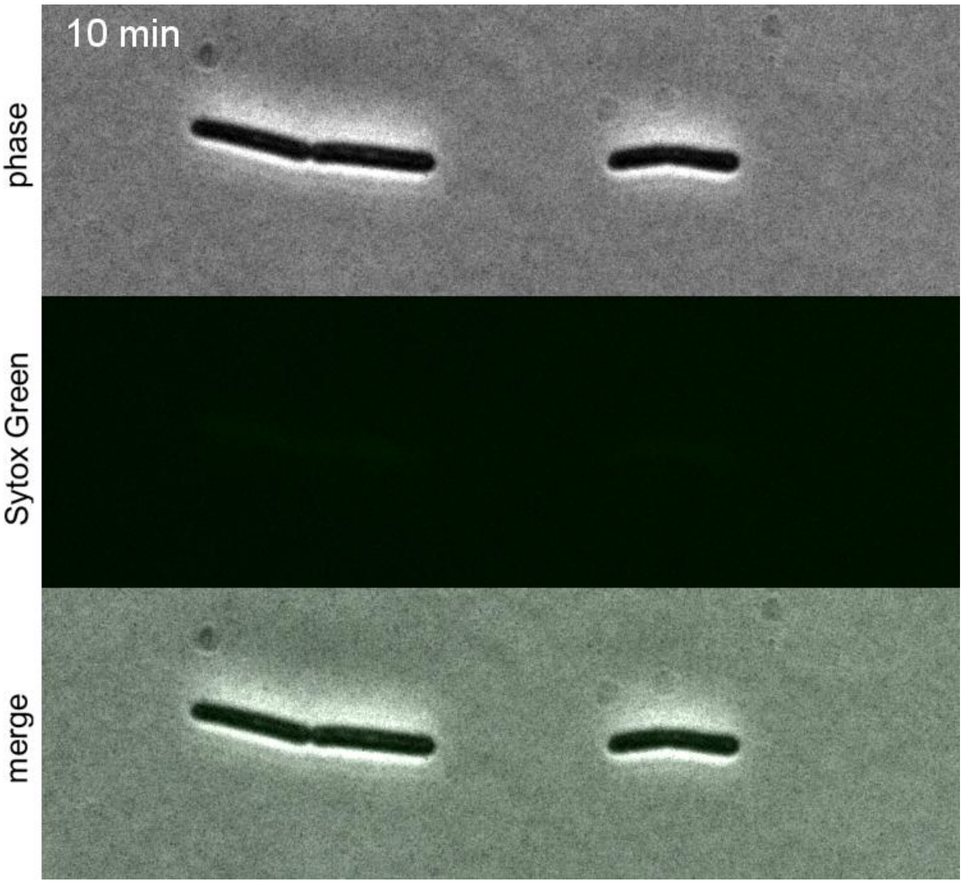


**Movie S1: Autolysis of *B. subtilis* can be monitored with the membrane permeability-indicator SYTOX Green**

*B. subtilis* phase contrast and fluorescence time-lapse microscopy of cells incubated on LB time-lapse slides and stained with 200 nM SYTOX Green in the presence of 100 µM of the proton uncoupler CCCP. Strain used: *B. subtilis* wild type (168).
